# Proteomic characterisation of matrix vesicles from primary osteoblasts indicates a mixed population with both exosomal and ectosomal routes of biogenesis

**DOI:** 10.1101/2025.11.07.687130

**Authors:** Scott Dillon, Charlotte Clews, Dominic Kurian, Fabio Nudelman, Colin Farquharson, Louise A. Stephen

## Abstract

Matrix Vesicles are a crucial step in producing a mineralised, healthy skeleton. Released from chondrocytes and osteoblasts, they concentrate calcium and phosphate to establish the deposition of hydroxyapatite around and within the collagen fibrils of the extracellular matrix, becoming embedded in the matrix in the process. In the 55 years since H. Clarke Anderson first described them, blebbing from the surface of chondrocytes, a consensus on their role, contents and their biogenesis, has yet to be reached. This is in part due to the range of cell types from which they are released and the multitude techniques that can be employed to isolate them. In this study, we, for the first time, characterise the proteome of matrix vesicles isolated from primary osteoblasts. By focussing on those vesicles that have become embedded within the matrix, we are able to avoid bias for specific modes of biogenesis and focus only on osteoblast-released vesicles that are associated with the matrix. Moreover, by studying these vesicles over a time course of mineralising activity, we are able to identify changes in the properties of these vesicles, and develop a more accurate picture of which proteins are involved specifically in mineralisation. In particular we identify the presence of markers associated with the ectosomal and exosomal release of matrix vesicles, as well as identifying proteins required for the maintenance and mineralisation of the extracellular matrix. These data portray a heterogeneous population of matrix vesicles, with different roles to play in bone development.

## Introduction

Matrix vesicle (MVs) biogenesis is a crucial but poorly understood initial step in physiological and pathological biomineralisation[1–3]. During biomineralisation, the extracellular matrix (ECM) of bone, cartilage, and teeth is strengthened by the deposition of mineral (poorly-crystalline substituted hydroxyapatite) around and within the collagen fibril [4, 5]. The protein-mineral composite is stiff and tough, providing bone with the flexibility and rigidity required for its structural roles in the skeleton, while conferring resistance to fracture. Moreover, the mineral component of the vertebrate skeleton is crucial for the maintenance of calcium-phosphorus homeostasis in the organism[6].

First identified in the late 1960s, MVs are small membrane-bound extracellular vesicles (EVs) of 50-200 nm [1, 7]. MVs act to concentrate calcium and inorganic phosphate, allowing for the transportation of calcium phosphate and eventual nucleation of crystalline mineral in the ECM [8]. While the importance of MVs for biomineralization is widely accepted, the process of their biogenesis and release from the cell is somewhat contested. Two pathways have been implicated; one in which vesicles ‘bleb’ from the plasma membrane of the osteoblast or chondrocyte, in an ectosome-like manner [1], and a second, recently observed pathway, reminiscent of exosome biogenesis, in which intracellular vesicles are transported to the membrane for release[9]. MVs have traditionally been studied as a distinct category of EV, containing hydroxyapatite [10] or precursor, amorphous calcium phosphate phases[11, 12], but other potentially critical vesicle contents have not been extensively studied. However, in recent years, several studies have identified additional MV properties that imply MVs may more closely resemble established EV types, such as exosomes (reviewed [13]). Like exosomes, MVs may also perform more complex functions through the transport of signalling or other bioactive molecules such as miRNAs [14, 15].

Proteomic analysis offers a key opportunity to document the composition of MVs, including information on how the vesicles have developed, how they are released, and what their contents and functions may be. Proteomics has been used to characterise MVs derived from mineralising cells *in vitro* and chicken embryonic femurs [16–19]. However, there are limitations to these studies. Thouverey studied osteoblast-like Saos-2 cells and isolated MVs from microvilli only, limiting any potential findings to the proteome associated with vesicles released via a microvilli-dependent ectosomal pathway[17]. Xiao and colleagues also used cell lines (MC3T3 osteoblast-like cells) to compare the proteome of MVs isolated from culture medium to those isolated from the ECM [19]. Balcerzak and colleagues isolated EVs from entire embryonic chick femurs, potentially capturing a very heterogeneous population of vesicles, including those released by non-mineralising cells [18].

Importantly, these studies employed qualitative proteomic methods, which prevented the analysis of protein abundance and changes in their expression over a mineralizing time course [17–20].

An important consideration in revisiting proteomic analysis of MVs is the recent implication of exosomal pathways in MV biogenesis [15, 21]. Earlier studies that focused on ectosomal pathways have risked biasing against these populations. For an in-depth review of potential pathways involved in MV biogenesis, please see [13]. In brief, ectosomal biogenesis is a process by which cargoes accumulate at the plasma membrane before being directly shed via budding. Exosomes, on the other hand, are characterised by intracellular packaging of cargo within intraluminal vesicles (ILVs). ILVs are, in turn, enveloped within a multivesicular body (MVB) that is transported via the endosomal pathway for transport to the plasma membrane, with which it fuses to allow release to the extracellular environment [13, 22].

Previous studies focussed on the proteomic composition of MVs have not been conducted within the guidelines of minimal information for studies of extracellular vesicles (MISEV) [23]. As we develop a more rounded understanding of MV biology, it becomes crucial to use these data in the context of broader EV research. With regards to MVs, it is pertinent to consider the conditions in which cells were maintained and isolated, in particular pH and temperature. By considering the MISEV guidelines in MV research, this study with primary murine osteoblasts reports a modified protocol for the isolation of a pure and complete population of biologically normal MVs. Proteomic characterisation of these vesicles allows for the further study of MVs within their EV context and supports the developing theory that MVs may be released through both ectosomal and exosomal pathways.

## Materials and methods

All materials were purchased from Sigma Aldrich unless otherwise specified.

### Collagenase-released matrix vesicle isolation

All animal studies were approved by the Roslin Institute’s named veterinary surgeon and named animal care and welfare officer, with animals maintained in accordance with the Home Office code of practice (for the housing and care of animals bred, supplied or used for scientific purposes). Animal studies were conducted in line with the ARRIVE guidelines.

C57BL/6 mouse pups of both sexes were sacrificed by the schedule one method of decapitation at 3-5 days old. Primary calvarial osteoblasts were isolated according to standard protocols [24]. Cells were plated at 200,000 cells in 100mm diameter cell culture-treated dishes in αMEM (Gibco, ThermoFisher Scientific, Waltham, USA) supplemented with 10% foetal bovine serum and 50µg/ml gentamicin at 37°C in a 5% CO_2_ atmosphere until 80% confluent, which was designated as day 0. Cells were stimulated with osteogenic media by the addition of 50µg/ml L-ascorbic acid and 5mM β-glycerophosphate or 10mM inorganic phosphate for up to 21 days, with media changed every 2 - 3 days. Mineralisation was assessed initially using phase contrast microscopy before alizarin red assays were performed according to standard protocols, in samples fixed in 4% paraformaldehyde for 10min at room temperature [24].

For collagenase-released matrix vesicle (CRMV) isolation, all solutions were filtered through a 0.1µm syringe filter to remove large contaminating particles and EVs isolated using differential ultracentrifugation as previously established [25]. Media was removed and the cell/matrix monolayer was washed thoroughly in Hank’s buffered salt solution (HBSS). The cell/matrix monolayers were solubilised by digestion in 2.5mg/ml collagenase IA in HBSS for 2 - 6h at 37°C with firm agitation until solubilisation was complete. Cells at more advanced stages of mineralisation requiring longer incubation. EVs were isolated by differential ultracentrifugation using the following protocol: centrifugation at 2,000xg at room temperature for 5min to pellet intact cells; supernatant centrifuged at 10,000xg at 4 °C for 30min to pellet cell debris; supernatant centrifuged at 100,000xg at 4 °C for 70min; supernatant discarded and EV pellet resuspended in 50mM Tris-HCl, pH 7.6. The solution was centrifuged at 100,000xg at 4 °C for 60 minutes, and the EV pellet was resuspended in 250µl 50mM Tris-HCl, pH 7.6, for transmission electron microscopy (TEM) and nanoparticle tracking analysis (NTA), or lysed in 20µl 8M urea, 0.1% SDS in 50mM HEPES, pH 8.0 for proteomics. Isolated MVs were snap frozen in liquid nitrogen and stored at -80 °C before analysis.

### Extracellular vesicle characterisation

NTA was performed using a Nanosight LM14 instrument (Malvern Instruments) equipped with a 532nm laser and a CMOS Hamamatsu Orca Flash 2.8 camera, operating with the NTA v2.3 software. Vesicle isolates were diluted 1:1000 in vesicle isolation buffer (50mM Tris-HCl buffer, pH 7.6), vortexed briefly, and particle concentration and size were measured using commercial algorithms. Measurements were repeated 3 times on each sample and the results were averaged.

For TEM, a drop of vesicle suspension was incubated on a 200-mesh formvar-coated copper grid (Agar Scientific, Rotherham, UK) for 10min. Excess solution was removed by touching the grid edge with filter paper, and vesicles were fixed in 2.5% paraformaldehyde, 2.5% glutaraldehyde in 0.1M sodium cacodylate for 2min. Grids were washed several times in distilled water and stained in 1% aqueous uranyl acetate (Agar Scientific). Grids were imaged using a Tecnai F20 field emission gun TEM (FEI), operating at 200keV and equipped with a CMOS camera.

### Quantitative proteomics

Total protein concentration was determined using the BCA assay (ThermoFisher Scientific, Altrincham, UK). Mass spectrometry was performed by the Roslin Institute Proteomics Facility. Samples were first reduced and alkylated, followed by digestion with trypsin. The resulting peptides were desalted by solid phase extraction on C18 cartridges (Affinisep, Normandy, France). The cleaned peptides were labelled with stable-isotopic iTRAQ 4-plex labels (Sciex Framingham, MA, USA) Following labelling, peptides were pooled and fractionated by strong cation exchange chromatography. Individual fractions were analysed by liquid chromatography-mass spectrometry, and datasets were combined to generate a protein list. Identification was performed using the Mascot software, and median ratios calculated using the ProteinScape (Bruker, Coventry, UK) and WARP LC (Bruker) software suite.

Analysis was repeated in 3 biological replicates, each consisting of cells pooled from different mice. Data were first filtered to exclude proteins identified using < 2 unique peptides. Data were expressed as a log_2_ fold change compared to Day 0 control samples.

### Data Availability

The proteomics data including raw files and search results have been deposited to the ProteomeXchange Consortium via the MassIVE partner repository with the dataset identifier PXD069400

### Bioinformatic analysis

Data analysis was performed in R. Gene Ontology (GO) enrichment analysis was performed using ClusterProfiler [26] and ECM annotation using MatrisomeDB and Matrisome AnalyzeR [27, 28].

For network analysis, processed data were imported to Graphia (https://graphia.app/) and subject to pairwise correlation analysis, with a cutoff value of < 0.97. Data were then subject to Markov cluster algorithm (MCL) clustering with a granularity value of 1.2 to generate clusters of proteins with correlated expression trends [29]. Protein-protein interaction networks were calculated using StringApp [30, 31]. A confidence score cut off of 0.9 was used and proteins with <2 predicted interactors were excluded.

## Results

### Osteoblast-derived collagenase-released matrix vesicles represent a unique population of extracellular vesicles

To induce the production of MVs, primary mouse calvarial osteoblasts were treated with osteogenic media supplemented with 10mM inorganic phosphate, which upregulates their release from mineralising cells as previously shown [25]. Phase contrast microscopy and alizarin red staining demonstrated the expected progression of matrix mineralisation over a 21-day post-confluence time course in osteogenic media (Fig. 1A). To ensure high phosphate treatment did not impact cell survival, we performed the alamarBlue cell viability assay in cells treated with 50µg/ml L-ascorbic acid only, conventional mineralising media conditions (50µg/ml L-ascorbic acid + 5mM β-glycerophosphate), and 10mM inorganic phosphate, revealing no significant differences in viability between conditions over the 21-day time course (Fig. 1B).

**Fig. 1.**
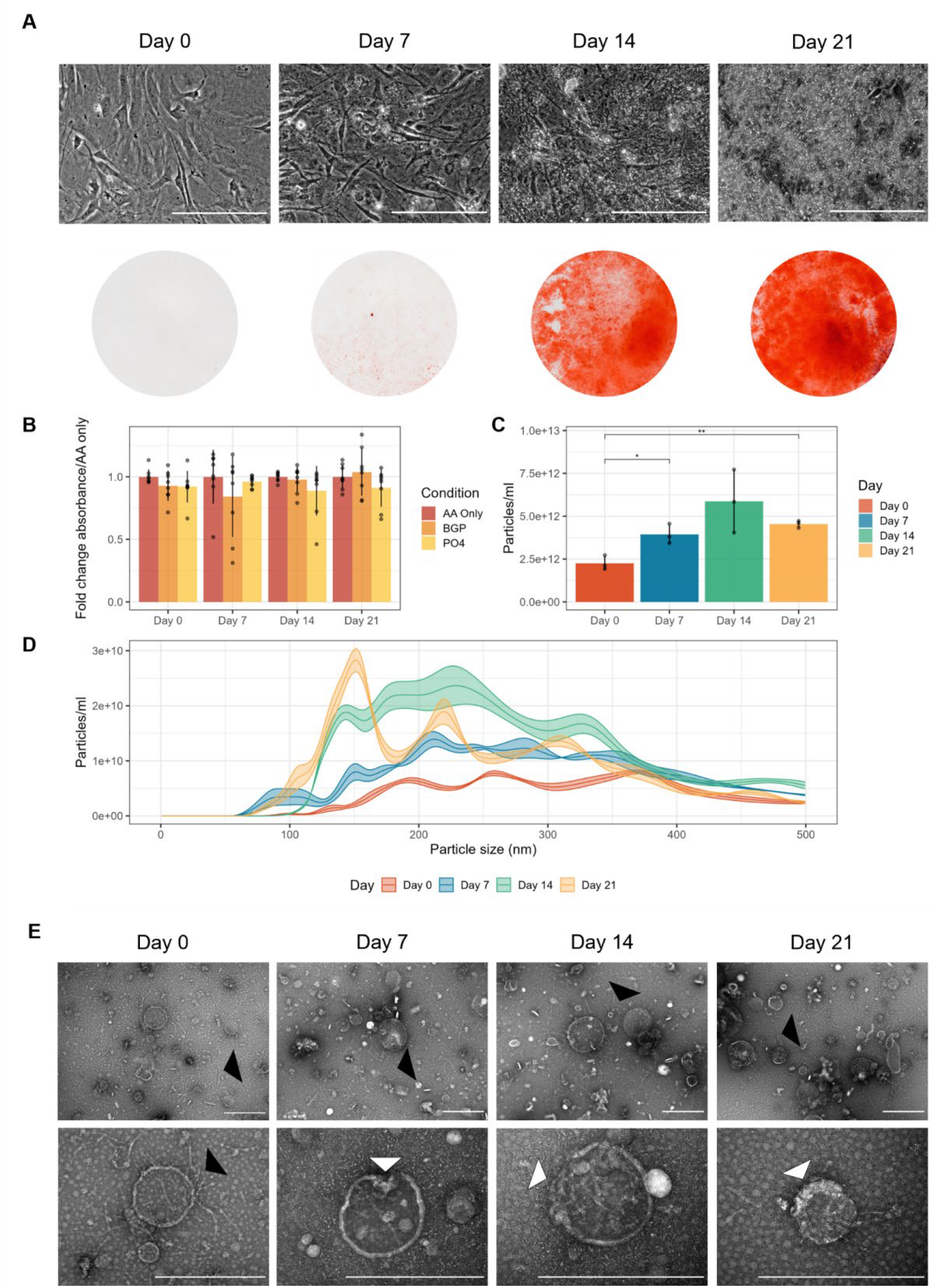
A) Phase contrast microscopy and alizarin red staining of primary calvarial mouse osteoblasts treated with osteogenic media supplemented with 10mM inorganic phosphate. Scale bars = 200µm. B) AlamarBlue cell viability assay over 21 days of culture showing no difference between treatments. C) NTA of CRMV isolates shows a significant increase in vesicle concentration over time. * p < 0.05; ** p < 0.01. D) NTA demonstrates a change in vesicle diameter over the mineralising time course. E) Negative stain transmission electron microscopy shows unilamellar vesicles present at all time points. Banded fibrils were often observed bound to vesicle surfaces (black arrowheads). Vesicles often harboured disrupted or crenelated membranes in Day 7, 14 and 21 samples (white arrowheads). Scale bars = 500nm

NTA of CRMVs demonstrated a significant (p < 0.01) increase in EV concentration in isolates from day 0 to day 14 in mineralising conditions (Fig. 1C). The distribution of vesicle diameter was also strongly regulated over time, changing from a broad relatively undifferentiated distribution between 50-500nm at 0 and 7 days of treatment, to defined peaks at day 21 with maxima at ∼150nm, ∼220nm and 305nm, suggesting enrichment of specific vesicle populations (Fig. 1D). Negative stain TEM was used to visualise isolated EVs (Fig. 1E). All samples were observed to contain unilamellar EVs along with banded fibrils 5-7.5nm in diameter which often associated with EV membranes (Fig. 1E, black arrowheads). These likely represent digested ECM fragments. Day 0 EVs appeared to have largely intact membranes in contrast to later time points, which exhibited many vesicles with disturbed or crenelated focal membrane regions (Fig. 1E, white arrowheads).

To define the protein composition of osteoblast-derived CRMVs, we performed quantitative proteomics on vesicle isolates. 1216 proteins were identified after exclusion of those classified using less than 2 unique peptides and with a large coefficient of variation. These comprised some classical osteoblast markers, including, for example, tissue nonspecific alkaline phosphatase (TNAP), osteopontin (OPN) and ectonucleotide pyrophosphatase/phosphodiesterase 1 (ENPP1) (Fig. 2A). Isolates were heterogeneous in cargo and displayed markers of both exosomes and ectosomes (Fig. 2A). Established exosomal markers included CD63, CD9 and tumor susceptibility gene 101 (TSG101). Those that are more characteristic of ectosomal pathways included integrins (such as integrins β1 and α5), CD81, and heat shock proteins (including heat shock protein 90α), but not CD40. Markers of earlier endosomal pathway steps were also present, including, for example lysosomal associated protein 1 (LAMP1). The isolated vesicles, therefore appeared heterogeneous, with potentially several populations present. Comparison of isolates with the EV proteome database ExoCarta revealed 467 (38.4%) shared proteins [32]. Functional enrichment analysis of 749 proteins unique to osteoblast-derived CRMVs revealed significant enrichment of GO Biological Process terms associated with cell-matrix adhesion, regulation of the actin cytoskeleton, and vesicle organization/transport (Fig. 2C). Similarly, significantly enriched GO Molecular Function terms included terms associated with actin binding, phospholipid binding, and adhesion/integrin molecule binding (Fig. 2D).

**Fig. 2.**
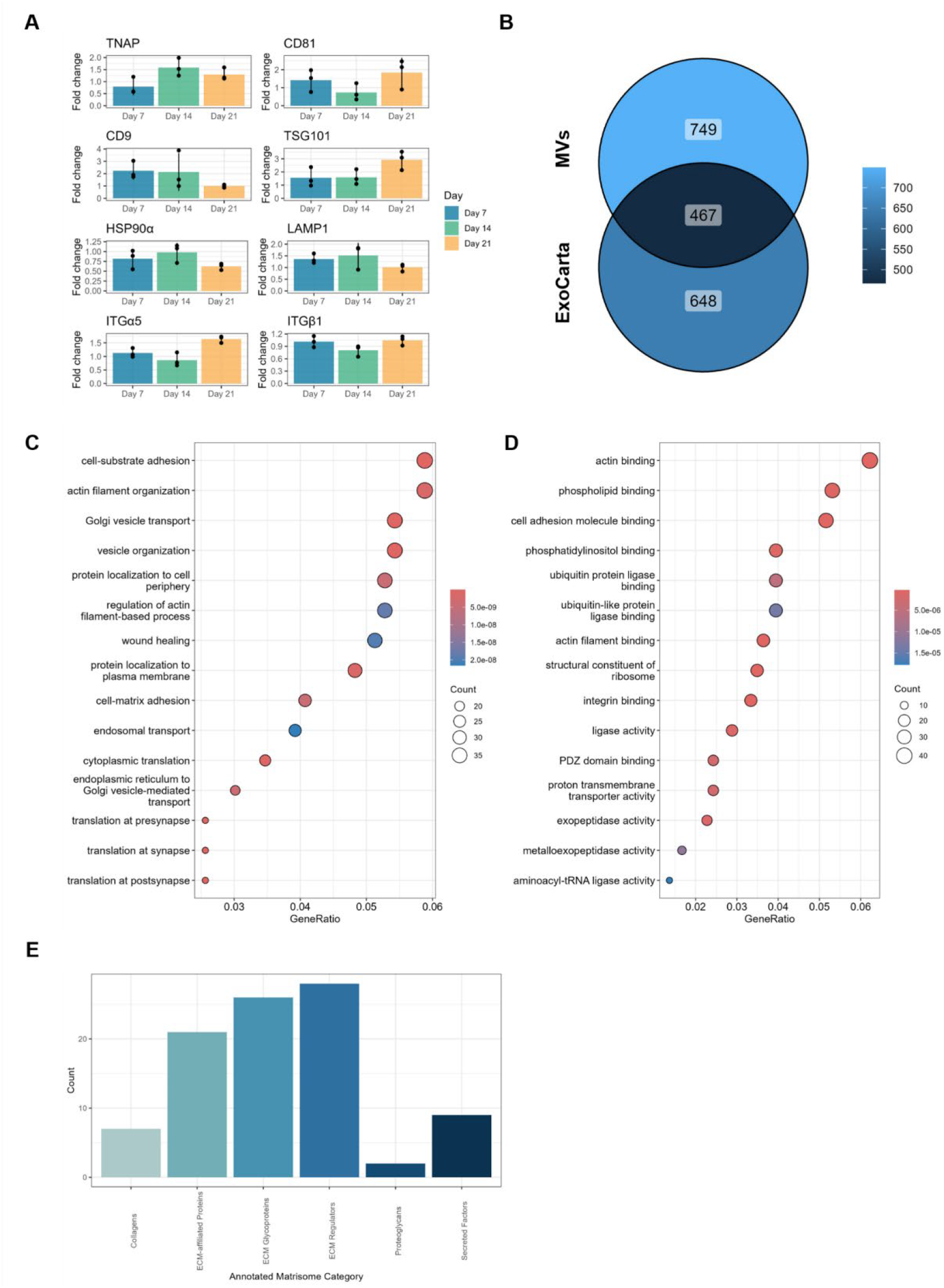
A) Quantitative proteomics data identify the presence of established osteogenic markers (e.g.) TNAP and markers of different EV populations. Data represent log_2_ fold change/Day 0 samples ± standard deviation. B) Venn diagram showing proportion of total CRMV dataset represented in the ExoCarta database. C,D) Functional Gene Ontology (GO) enrichment analysis performed on the total dataset for Biological Process (C) and Molecular Function (D) terms. E) Annotated extracellular matrisome categories of identified proteins

Unique CRMV proteins were compared against the ECM proteome using MatrisomeDB (Fig.2E) [27, 28]. Several structural ECM molecules were identified, including α chains of collagens V, VI, XII, and XVIII, fibronectin, and laminin β1. Several ECM proteases were also identified, including matrix metalloproteases (MMPs) 2 and 14 and several members of the disintegrin and metalloproteinase domain-containing protein family (ADAM9, 10, and 17, and ADAMTSL4). CRMVs therefore, appear to be strongly associated with structural components of the ECM and may actively modify the matrix within which they are embedded.

### Time-resolved network analysis disentangles vesicle functional populations

To separate heterogeneous vesicle populations we performed network analysis of the quantitative proteomics data, using time under mineralising conditions as a covariate. Graphia was used to calculate and visualise the pairwise correlation matrix between identified proteins using a correlation score cut-off of 0.97 (Fig. 3A) [33]. MCL clustering was used to generate clusters of proteins with correlated expression over time, identifying 5 clusters in total (Fig.3A-C).

**Fig. 3.**
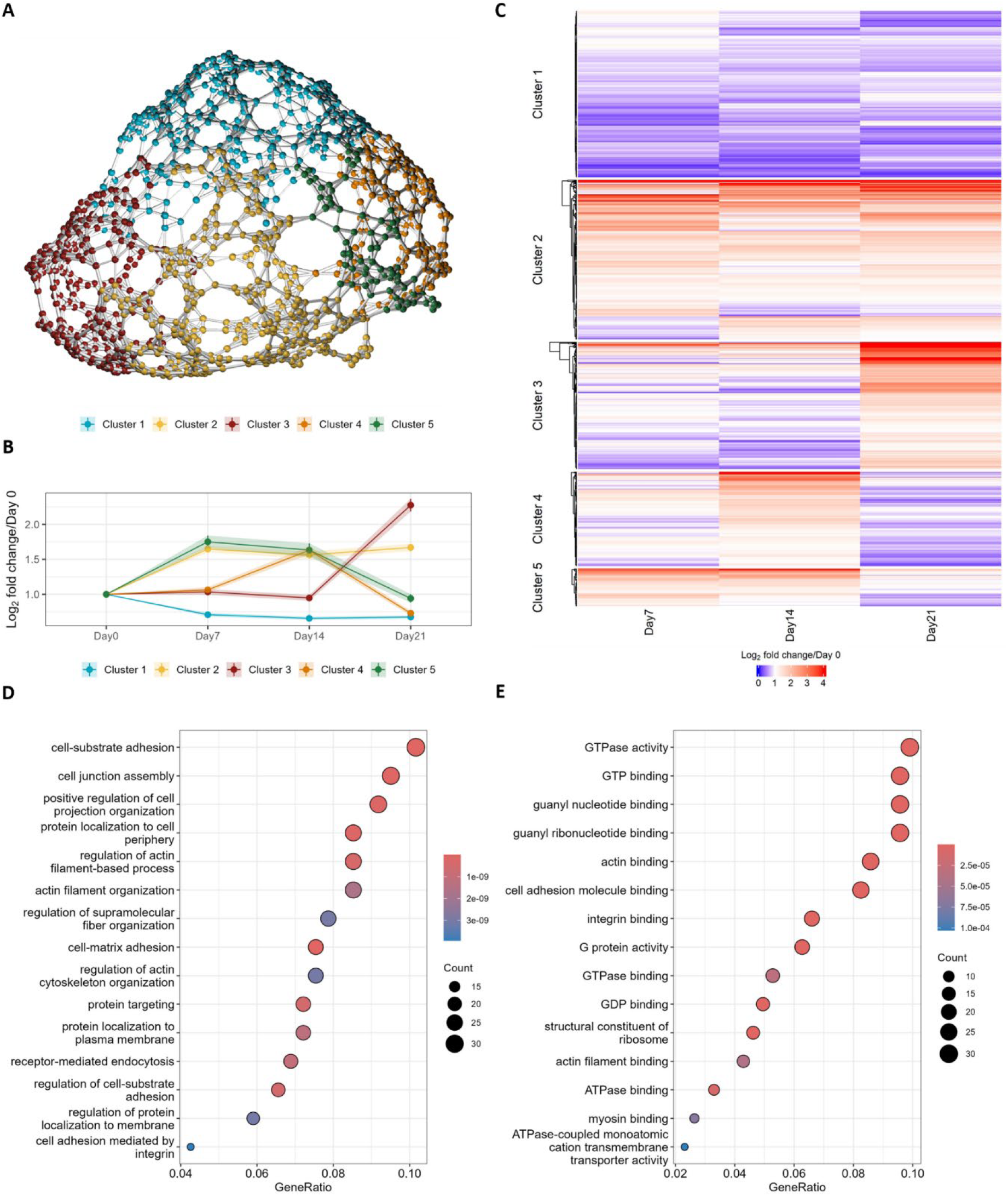
A) Total pairwise correlation matrix network, visualised using Graphia. Nodes represent individual proteins and edges represent correlation of expression. Nodes were separated into 5 clusters based on expression over time using MCL clustering. B) Average expression over time of proteins in each cluster. Data represent mean log_2_ fold change / Day 0 samples ± standard error of the mean. C) Heatmap showing absolute log_2_ fold change / Day 0 expression of all identified proteins grouped by cluster. D) GO enrichment analysis performed in Cluster 2 proteins for Biological Process (D) and Molecular Function (E) terms

Temporal protein expression patterns are likely related to protein function and the phenotypic status of the cells in culture. Upon stimulation with osteogenic media primary osteoblasts appear to downregulate cluster 1 proteins within their extracellular vesicle cargo and therefore these are unlikely to be directly related to MV-mediated biomineralisation (Fig. 3B). Conversely, cluster 2 proteins are actively upregulated on average and therefore are more likely to have been actively trafficked to vesicles to mediate biomineralisation (Fig. 3B). All major osteogenic markers identified including TNAP, OPN and ENPP1 fall within cluster 2.

GO functional enrichment analysis was performed on Cluster 2 proteins, revealing a significant enrichment of Biological Process terms associated with cell-ECM adhesion and the actin cytoskeleton (Fig. 3C). Significantly overrepresented Molecular Function GO terms demonstrated enrichment of both actin binding proteins and G proteins/GTPases (Fig. 3D). A large proportion of these proteins had functions involving regulation of actin dynamics at the terminal ends of filaments, including cofilin-1, profilin 1, formin-like 2, tropomodulin-3 and cortactin. Among the variety of GTPases identified were Rab vesicle trafficking proteins, along with associated G proteins. Interestingly, Ras homolog family member A (RhoA) and RAS-related C3 botulinum substrate 1 (Rac1) and cell division control protein 42 (CDC42) which are major antagonistic GTPases regulating cytoskeletal dynamics were also present [34].

Clusters 3-5 include proteins that exhibit variable average expression patterns over the course of mineralisation, suggesting that these may be associated with mineralisation, but may also correlate with other cellular processes. Within these clusters are key proteins associated with exosome biogenesis, in particular, syntenin and syndecan1 and 2 and others associated with cytoskeletal dynamics (e.g., spectrin beta, non-erythrocytic 1 (STBN1), and capZ alpha and beta subunits; Cluster 3)[35].

Collectively these data indicate that proteins regulating cytoskeletal dynamics are upregulated in CRMVs over a mineralising time course and may implicate the actin cytoskeleton in regulation of MV biogenesis.

### Protein-protein interaction networks implicate cytoskeletal dynamics in vesicle biogenesis

To elucidate functional relationships between cluster 2 proteins thought to be involved in cytoskeletal regulation of MV biogenesis, protein-protein interactions were visualised using StringApp (*5, 6*). 267 of 301 proteins were found to have more than 1 interaction (Fig. 4A). The network exhibited several dense interconnected regions, including one centred around Rac1, RhoA and CDC42 (Fig. 4B). These regulatory GTPases interacted directly with cluster 2 proteins regulating actin severing and depolymerisation, such as cofilin-1, profilin-1, cyclase-associated protein 1 (CAP1) and destrin, and polymerisation of branched actin networks such as components of the actin-related protein (ARP) 2/3 complex (Fig. 4B).

**Fig. 4.**
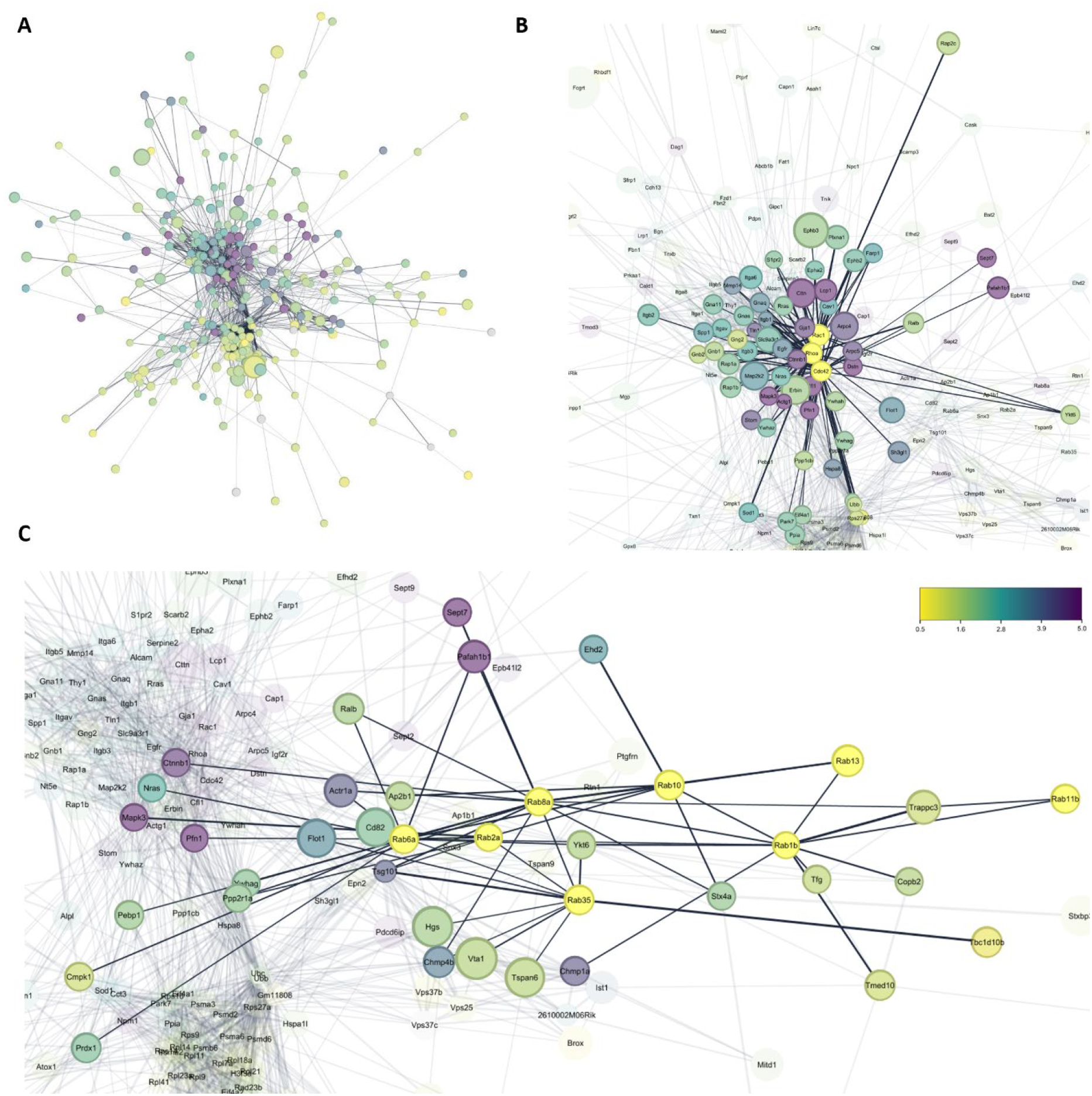
Protein-protein interaction network calculated from cluster 2 proteins, excluding nodes with < 2 interactions (34/301 total). A) Total calculated network. B) Direct interactors of the cytoskeleton regulating GTPases RhoA, Rac1 and CDC42. C) Direct interactors of RAB vesicle trafficking proteins identified in cluster 2

A distinct region of the network was enriched in RAB vesicle trafficking proteins, including RAB1b, 2a, 6a, 8a, 10, 11b, 13, and 35, several of which are implicated in transport between the Golgi apparatus and endoplasmic reticulum or in endocytic recycling (Fig. 4C) [36, 37]. Several of these components interacted with the actin-regulating region through mitogen-activated protein kinase 3 (MAPK3), profillin-1, the GTPase-activating protein NRas, or β-catenin (Fig. 4C).

Functional analysis of protein interactions in CRMVs therefore suggests that vesicle cargo is regulated by endosomal trafficking, and that intracellular transport or EV biogenesis is governed by dynamic regulation of actin severing and depolymerization through RhoA/Rac1/CDC42 signaling.

## Discussion

MVs present an exciting front in understanding the cellular control of biomineralisation and bone development, and as such have attracted wide-ranging applications to their study. This has led to opposing strategies, some of which fail to take advantage of modern technologies. Whilst the use of cell lines as described in previous proteomic analyses of MVs, can offer a more homogenous, well-characterized route to analysis, they risk introducing non-physiological EV characteristics due to the genomic changes identified in immortalised cells such as MC3T3 and Saos-2 cells [17, 19, 20]. This paper represents the first use of primary osteoblast cells to characterise the MV proteome. In the current study, MVs were isolated from a digested mineralising ECM to ensure the study specifically targeted MVs rather than including other, non-MV, EVs released from osteoblasts into the culture medium [38, 39]. By investigating CRMVs, we are better placed to avoid the pitfall of misclassifying EVs based on their expected role or biogenesis pathway. We have also quantified temporal expression patterns in the MV proteome over a mineralisation time course for the first time.

The CRMVs isolated from primary osteoblast cultures appear physiologically normal, with intact unilamellar membranes at day 0 of mineralisation, with increasing membrane disruption at day 7, 14 and 21 (Figure 1). These data support the degradation of the vesicle phospholipid membrane as a source of extracellular inorganic phosphate as previously proposed [24, 40–43]. It is notable that only 38 % of proteins identified in our MV populations are also identified in the ExoCarta sEV protein database [32]. There are likely two reasons for this, firstly, MVs appear to be specialised EVs that contain proteins not specifically associated with traditional EV populations. Some of these are identified in Figure 2E, in which we identify 98 proteins from our screen that are associated with the ECM. However, CRMVs are also likely to have undergone unique biological changes to allow for the deposition of calcium and phosphate for the initiation of biomineralisation. These MVs are beginning to break down their membranes, to allow for the release of their contents. The exact process by which membranes breakdown is not known, however, two modes are hypothesised; Firstly, that membranes are ruptured by crystalline HA as it penetrates the membrane, or secondly, that the MV membrane is weakened by the breaking down of the phospholipids such as phosphatidylcholine and phosphatidylethanolamine within the MV membrane during the mineralisation process [40, 44]. Whatever the process, these alterations will likely result in changes to the proteomic profile of the vesicle. We must therefore be clear in our interpretation of these data that we are examining all vesicles that have been released from osteoblasts and have become embedded in the mineralised collagen matrix of these cultures. This may mean that vesicles, not yet embedded within the ECM, but with the potential to mineralise collagen are not captured. It is also possible that those vesicles that are captured may not be solely those traditionally considered ‘matrix vesicles’.

By conducting a temporal analysis of CRMV proteins from non-mineralising through to established stages of mineralisation, it is possible to identify the proteins required for the release and function of MVs, without being limited by any pre-existing assumptions. The increase in vesicles isolated across the expected timeline of mineralisation and their enrichment for osteogenic markers allows a strong level of confidence that these are MVs. However, there are limitations to the study. Along with the considerations noted above with regard to post-embedding biochemical changes, key markers are absent from our data. Most prominently perhaps, the lack of PHOSPHO1. This phosphatase is known to be present in MVs and is considered a vital step in the enrichment of MVs for phosphate [45, 46]. Likewise, while Syntenin is recognised as the key marker of exosomal identity, its partner protein, ALIX, would similarly be expected to be identified in any exosomal population; its absence from this data set is unexpected [35]. However, these are proteins associated with the production, transport, and enrichment of vesicles prior to becoming embedded within the ECM. This analysis may highlight the dynamic nature of vesicles throughout their lifetime, and once embedded within an ECM, MVs have no requirement for PHOSPHO1 and ALIX.

The language around the release of vesicles is much debated, and researchers should refer to the MISEV guidelines for guidance [23]. Whatever the biogenesis pathway of MVs, the process will rely on the rearrangement of the cytoskeleton, as identified in our data [1, 15, 20, 47]. The data presented here show evidence of a heterogeneous population, in which vesicles are released via both an ectosomal and exosomal pathway. This is in support of recent work in which Boonrungsiman and colleagues produced novel EM data in which ectosomal release was identified alongside MVBs enriched with calcium [21]. Likewise, a recent paper from Binet and colleagues identify two subpopulations in the total EV load released from bone tissue [48]. The Rac1 cluster, within cluster 2 (Figure 4 B), consistently upregulated with the onset of mineralisation, is associated with the formation of microvilli, potentially supporting the microvilli, ectosomal hypothesis [20]. While CD81 is often cited as an exosomal marker, Mathieu and colleagues suggest it is specific to a subpopulation of ectosomes, its presence in cluster 2 adding to the identified ectosomal markers [49]. However, the identification of Rab7 and Rab8, alongside the endosomal sorting complexes required for transport (ESCRT) proteins in cluster 2, suggests the involvement of the endosomal pathways required for exosomal biogenesis. Moreover, Syntenin1 is considered the exemplar marker of exosomes [35]. Its presence, and that of Syndecan1 in cluster 3 of our samples, alongside CD63, confirms that within the CRMV population, are certainly a population of MVs that are produced via MVBs, i.e., exosomes [35, 49]. Figure 5 presents a schematic of key proteins identified within this dataset and their role in MV biology.

**Fig. 5.**
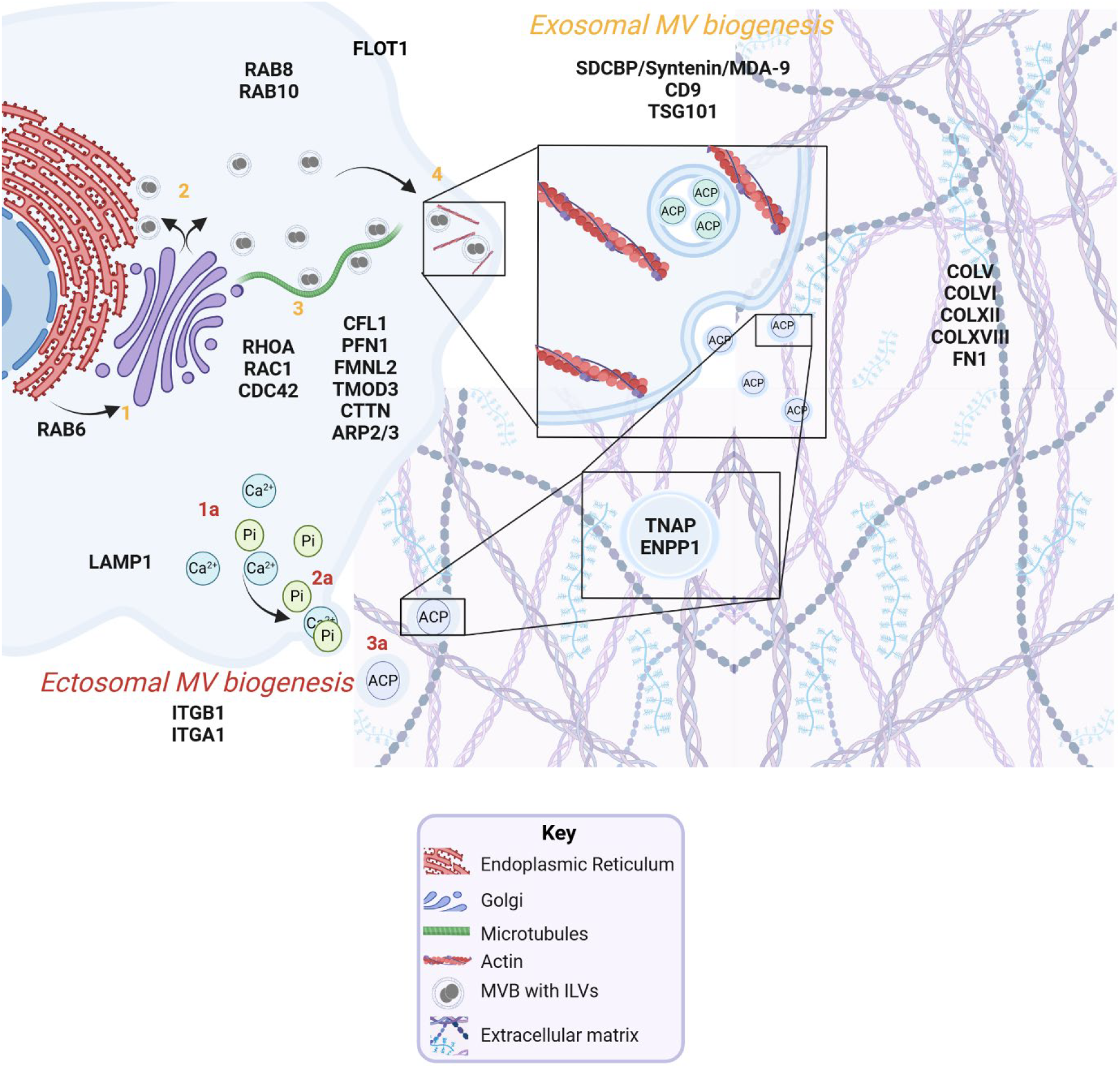
Proteomic analysis of CRMVs identifies a heterogenous population of vesicles released via both exosomal (1 – 4) and ectosomal (1a – 3a) pathways. Proteins associated with each stage are shown near their corresponding sites of action (black text). Exosomal MV biogenesis: After protein processing in the endoplasmic reticulum and Golgi apparatus (1–2), calcium phosphate containing ILVs are enclosed within intracellular MVBs. These MVBs travel along the cytoskeleton (3) and subsequently fuse with the plasma membrane, releasing ILVs as MVs into the ECM. Ectosomal biogenesis: Calcium and phosphate ions first accumulate at the plasma membrane (1). A portion of the membrane then blebs, forming a calcium phosphate containing ectosome (2), which is released into the ECM as an MV (3). ACP=amorphous calcium phosphate; MVB=multivesicular body; ILV=intracellular vesicle

In addition to the insights into the biogenesis of MVs garnered in this new analysis, we are further able to develop our understanding of MV cargoes. Until recently, little attention has been paid to the non-mineral content of MVs, but recent studies have been developing the idea that exosomal properties of MV biogenesis accompany a broader cargo, more akin to that of exosomes [15, 50, 51]. While Skelton, Lin, and colleagues focus on miRNA cargoes, our proteomic analysis suggests protein cargoes linked to the maintenance of the ECM, are present. It is known that EVs are involved in this process, and therefore these data pose the question as to what constitutes a MV as opposed to an EV found within the collagenous matrix. While it is not possible to answer this question within the confines of this work, it seems likely that the vesicles studied here, which increase in number with mineralisation, are playing a role in ECM maintenance as well as mineralisation.

## Conclusions

The data presented here suggest a unique, heterogeneous population of EVs released by osteoblasts that are embedded within the ECM during collagen fibril mineralisation. This supports the hypothesis that there are two populations of MV, one secreted via the ectosomal route and another via the exosomal route (summarised in Figure 5). Furthermore, we present evidence that MVs are possibly capable of roles in addition to biomineralisation, and likely to be involved in ECM maintenance, suggesting a broader role for MVs than has been previously posited. Ongoing work should aim to determine whether these are distinct populations with defined roles, or the result of osteoblasts utilising every opportunity to release MVs and mineralise their matrix when stimulated to do so.

## Acknowledgements

The authors would like to thank Biotechnology and Biological Sciences Research Council (BBSRC) for supporting LAS via Discovery Fellowship (BB/X009904/1), SD through an EASTBIO Doctoral Training and Partnership studentship award (1803936) and for Institute Strategic Programme Grant Funding BBS/E/RL/230001C to the Roslin Institute which supports CF & DK.CC is gratefully funded by a PhD scholarship received from The University Edinburgh Doctoral College. For the purpose of open access, the authors have applied a CC-BY public copyright license to any Author Accepted Manuscript version arising from this submission.

## Author Contributions

Conceptualization: SD, CF, FN, LAS; Methodology: SD, CF, FN, DK, LAS, Formal analysis and investigation: All authors, Writing - original draft preparation: LAS, SD; Writing - review and editing: All authors, Funding acquisition: SD, CF, FN, LAS; Supervision: CF, FN, LAS

